# Transcriptomic Analysis of the Neurogenesis Signature suggests Continued but Minimal Neurogenesis in the Adult Human Hippocampus

**DOI:** 10.1101/664995

**Authors:** Ashutosh Kumar, Vikas Pareek, Muneeb A. Faiq, Pavan Kumar, Chiman Kumari, Himanshu N. Singh, Sanjib K. Ghosh

**Author notes:** Contributed equally (Ashutosh Kumar, Vikas Pareek). **Corresponding author** Dr. Ashutosh Kumar.

## Abstract

**Purpose:** Since immunohistological investigations have given rise to divergent perspectives about continued hippocampal neurogenesis in adult humans, a comprehensive transcriptomic analysis of the neurogenesis signature markers supplemented with insights from gliogenesis and apoptotic markers (in context to the developmental stages across age) may discern important aspects and may well be the appropriate methodology for resolving this conflict.

**Materials and Methods:** RNA expression data for the salient neurogenesis, gliogenesis, and apoptotic marker genes in post-mortem human hippocampal tissue of the Prenatal (n=15), Infant/Early childhood (n= 5), Adolescence (n=4), and Adulthood (n=6) ages were downloaded from Allen Brain Atlas database (http://www.brainspan.org/rnaseq). Gene expression data was categorized, median values were computed, and age group specific differential expression was statistically analyzed (the confidence level of 95%, p value ≤ 0.05 is used).

**Results:** A sharp fall in prenatal to infant/early childhood expression was observed for all studied neurogenesis markers, except that for the post-mitotic late maturation (CALB1, CALB2, MAP2, NEUN, STMN2) which showed no significant differences in expression profiles. A continued post childhood decrease across advancing age was observed in the neural stem cells and progenitor markers with insignificant differences across close age groups. Uniquely, the postnatal sharp fall of KI67 and TBR2 continued across advancing age groups, reached near baseline until adolescent age. The immature granule cell, post mitotic early maturation, and late maturation markers showed a maintained or slightly increased (albeit insignificant) post childhood expression. The gliogenesis markers mostly showed a significant downregulation between prenatal and infant /early childhood age groups; this decline was persistent thereafter with insignificant differences between close age groups. A continued postnatal decrease occurred in apoptotic markers with observable, but insignificant, differences between adolescent and adult age.

**Conclusions:** Our findings indicate a possibility of continued but minimal neurogenesis in the adult human hippocampus. A significant part of the newborn cells in the neurogenic niche may be glial cells.

**(Findings of this study were first presented at the Annual Meeting of Society for Neuroscience (SFN), 3^rd^-7^th^ November, 2018, San Diego, USA.** https://abstractsonline.com/pp8/#!/4649/presentation/37213)

Graphical Abstract

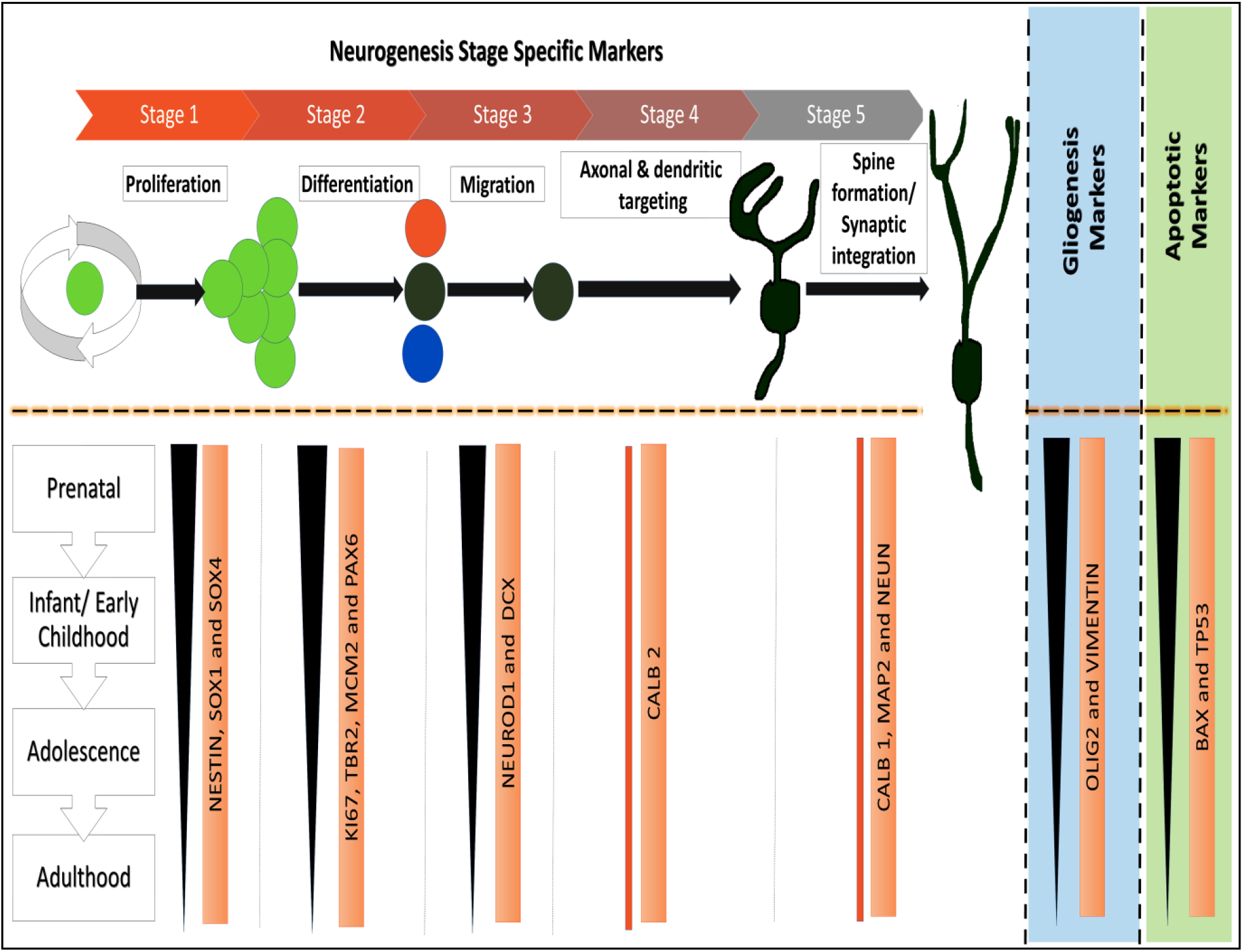

**Highlights:**

- A varying but continued fall in expression was observed for all studied neurogenesis markers across advancing age groups, except that for the immature granule cells, early and post-mitotic late neuronal maturation markers.
- A continued postnatal sharp fall of progenitor markers KI67 and TBR2 reached near baseline until adolescent age.
- Gliogenesis markers showed continued but insignificant fall in expression post infant/early childhood.
- Apoptotic markers showed continued post infant/early childhood downfall but changes were negligible between adolescent and adult age.

## Introduction

Continued adult hippocampal neurogenesis (AHN) in humans was recently questioned (1,2). Studies with contrasting evidence have emerged since then which ranged from absolute denial (3–5) to approval for the persistence of AHN even in old individuals (6–8). In adult mammals (including human) neurogenesis is thought to persist in the subgranular zone (SGZ) of the dentate gyrus (DG) region of the hippocampus (9). Newly formed neurons from the SGZ travel to hippocampal and prefrontal cortex (10), and integrate into the existing circuitries (10,11). Induced adult hippocampal neurogenesis was shown to improve spatial memory and performance at the learning tasks (12) in addition to be protective role against neuropsychiatric disorders which presumably precipitate a decline of these functions (12,13).

With the accumulation of optimistic evidence, AHN in most of the mammals (including humans) grew into a dogma (1,14,15). Dissenting results regarding AHN in any higher order mammal was rare (16); though in recent years, a few studies have denied AHN in human (1). Dennis et al, 2016 denied AHN in humans using immunohistological methods in postmortem samples across different age groups (4). The most robust disapproval, however, has appeared recently from Sorrells et al (3), and it has been followed by a similar denial from Cipriani et al (5), who based their conclusions on a more comprehensive use of immunohistological and ultrastructural methods, and in a greater diversity of samples. All three AHN negative studies shared the view that if any newly formed cells in the neurogenic niche are found, those are actually glial cells. The denial, nevertheless, was not long lived as AHN in humans was reaffirmed by studies and arguments from Boldrini et al (6). These investigators used similar immunohistological methods as were emplyed by Dennis group and Cipriani group., The Boldrini group supplemented their methodology by use of additional techniques thus employing a more convincing method of cell counting—unbiased stereology, measured accompanied vasculogenesis, and ensured that studied subjects didn’t suffer from any confounding disease before death (17).

More recently, more robust evidence emerged for AHN in humans (7,8). Moreno-Jiménez et al, 2019 using tightly controlled conditions and state-of-the-art tissue processing methods, identified thousands of immature neurons, exhibiting variable degrees of maturation along differentiation stages, in DG of the neurologically healthy human subjects up to the ninth decade of life. They additionally found that continued neurogenesis was reduced in DG of Alzheimer’s patients (8). Tobin et al, 2019 found AHN to persistent through the tenth decade of life and being detectable in patients with mild cognitive impairments and Alzheimer’s disease. These researchers found a reduction of AHN in mild cognitive impairments, and higher AHN associated with better cognitive status (8).

Hippocampal neurogenesis is believed to be a unique advancement in mammalian evolution though a protracted postnatal neurogenesis has been undoubtedly established in most of the studied mammalian species, with a few exceptions (18,19). JS Snyder, 2019, by drawing upon the published data on hippocampal neurogenesis across life span in most commonly studied mammals, observed that a lower rate of neurogenesis with accelerated neurodevelopmental timing is aligned across species (including humans) (20). New-born neurons during the protracted neurogenesis retain unique plastic properties for long intervals, and have distinct functions depending on when in the lifespan they were born (20). Based on these confirmations and arguments, continued formation of new neurons seems essential to accommodate new memories and provide adaptive flexibility to new life experience, hence present a survival benefit (20). A comprehensive review of the literature suggests that studies contesting for AHN in human stand in an arbitrary zone where even a minor difference of the investigatory methods and study designs may lead to alternative interpretations. Neurogenesis involves multiple developmental stages which are characterized by expression of specific protein markers which can be immunostained to observe the lineage specific cells in the neurogenic niche (21). Until now, these markers were mostly studied, in varying combinations by immunohistological methods within limited age groups, and (most importantly) their comprehensive transcriptomic analysis was largely ignored. A transcriptomic analysis of the differential expression patterns of neurogenesis signature markers, added with the gliogenesis and apoptotic markers (respective to the stages of neuronal maturation in developing to adult age hippocampus) may show or construct the much needed larger picture. It may also resolve the apparent divergence in the interpretations from immunohistological studies. Though this method cannot provide an accurate quantitative estimate of the newly formed neurons (or glial cells), it can distinctively show the trends for the expression of the marker genes in advancing age groups, which can be inferred for the perpetuation of AHN. With all this background, we analyzed the developmental transcriptome (prenatal to adult age) in human hippocampus to check the differential expression of the neurogenesis signature genes (and also gliogenesis and apoptotic markers) respective to the neuronal developmental stages to identify potential answers to the question of AHN.

## Materials and Methods

RNA expression data for the neurogenesis signature genes in post-mortem human brain tissue of the Prenatal (n=15), Infant-early childhood (up to 3 years age, n= 5), Adolescence (11-19 years, n= 4), and Adulthood (20-40 years, n=6) ages from the hippocampus were downloaded from development transcriptome database of Allen Brain Atlas (http://www.brainspan.org/rnaseq). Post-mortem brain specimens only from neurologically healthy individuals and free of significant genetic errors were considered for original data retrieval (to ensure consistency between samples and to decrease potential variation arising of ante- and postmortem conditions, specific selection criteria were followed, details for which, along with the protocols for the laboratory procedures and techniques used, can be found on Allen Brain Atlas website link: http://help.brainmap.org/display/devhumanbrain/Documentation?preview=/3506181/6651924/Transcriptome_Profiling.pdf).

Gene expression data was categorized as per age groups and median expression values were computed for the targeted neurogenesis signature genes (Table 1 presents the list of the neurogenesis stage specific signature genes selected for this study). In addition to neurogenesis signature genes, gliogenesis (OLIG 2, VIMENTIN, and S100B), and apoptotic marker genes (BAX and TP53) were also studied. Differential gene expression respective to the neurogenesis maturation stages across the studied age groups (Table1) was analyzed using statistical tests (differences in the median values with standard deviations). We used non-parametric statistical tests (owing to the unequal number of samples in each category). Kruskal-Wallis (KW) (equivalent of parametric One-way ANOVA) and Mann-Whitney U (MWU) tests were used to check if variations existed between all four age groups, and between any two age groups, respectively (Table S1&2). For both the statistical tests, the confidence level of 95% (p value <= 0.05) was used.

**Table 1.** Developmental stage specific expression of immuno-histological protein markers in neurogenic niche of adult hippocampus.

| Stage 1<br>(Proliferation/Neural stem cell markers) | Stage 2<br>(Differentiation/Progenitor intermediate cell forms markers ) | Stage 3<br>(Migration/Immature granule cell markers) | Stage 4<br>(Axonal & dendritic targeting/Early maturation markers) | Stage 5<br>(Synaptic integration/Late maturation markers) |
| --- | --- | --- | --- | --- |
| NESTIN, BLBP, GFAP, SOX1, 2 & 4 | KI67, TBR2, MCM2, PAX6 | NEUROD1, DCX*, PSA NCAM | STMN2, SEMA3C, Calretinin** (CALB2), TUBB3 | MAP2 Calbindin (CALB1), NEUN,*** |

\*Also expressed in Stage 4, \*\* Also expressed in Stage 5, \*\*\* Also expressed in stage 4

A post-hoc ‘Bonferroni Correction’ was applied to check errors caused due to multiple comparisons performed for same sample in MWU test. Box plots with median values and full range of subjective data (were plotted to get trends for the age group specific gene expression (Fig.1-7, S1-7).

## Results

(Table S1&2, Figure 1-7, S1-7)

**Figure 1.**
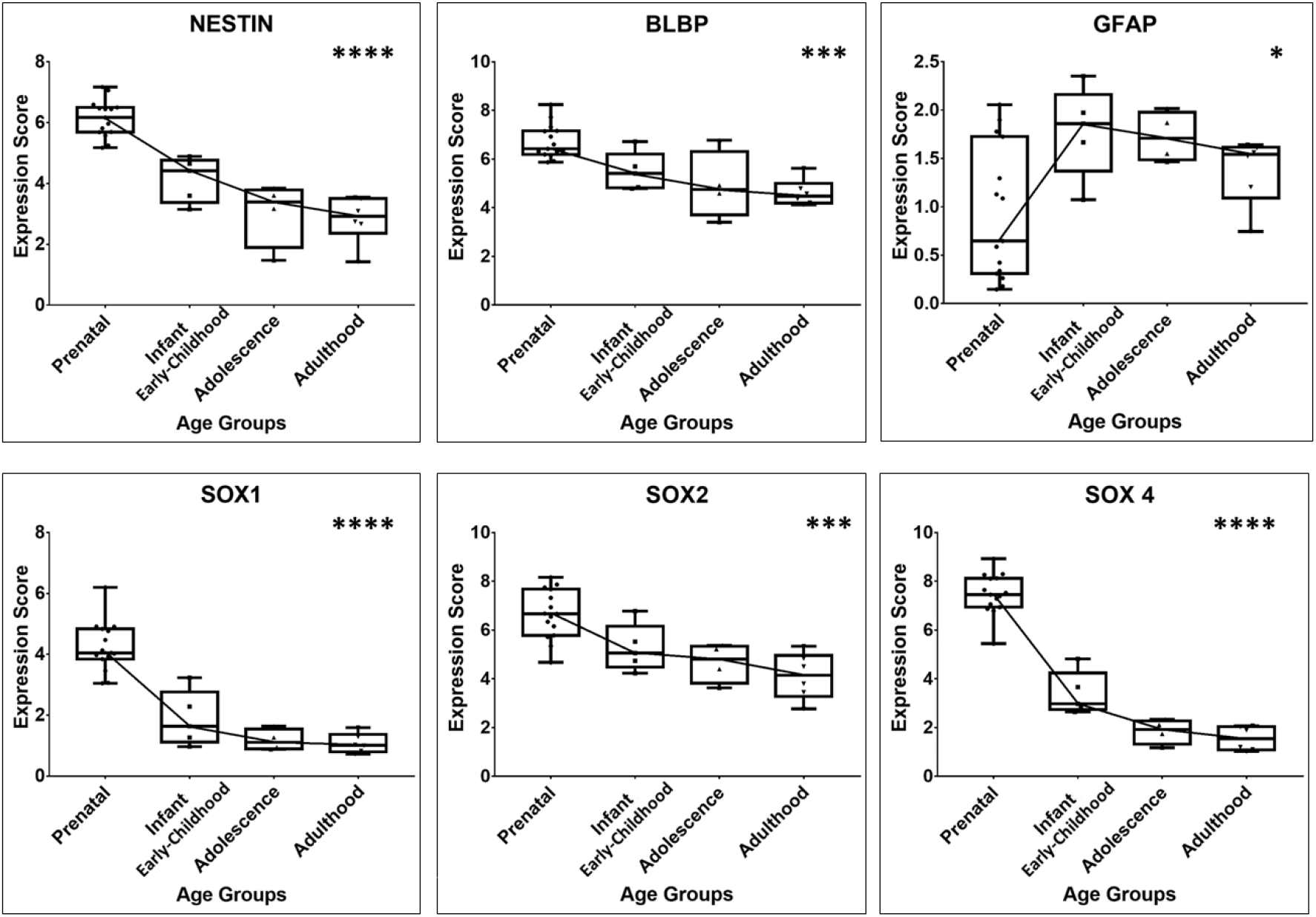
Box plot presentation of Neurogenesis Stage 1 specific gene expression scores (Statistical significance tested using Kruskal Wallis test, p values: *≤0.05, **≤0.01, ***≤0.001, ****≤0.0001.).

### Neurogenesis signature markers (Stage 1-5)

A prenatal to infant/early childhood sharp decrease in expressions of stem cell, proliferation, immature granule cell, and post mitotic early maturation, was noted with few exceptions (BLBP, GFAP, SOX2, PSA NCAM, CALB2, MAP2, CALB1, NEUN, all showing no significant differences) (Fig. 1-5). BLBP, and SOX2 (stage 1) didn’t show significant prenatal to infant/early childhood decline (Fig. 1, S1). GFAP (stage 1) showed insignificant increase between prenatal to infant/early childhood (Fig. 1, S1 (c.1)).

A post infant/early childhood continued decreasing trend across further age groups in expression occurred for the markers for stem cells (NESTIN, SOX1, SOX4)—Stage 1 (Fig. 1), and progenitor cells (KI67, TBR2, MCM2, PAX6)— Stage 2 (Fig. 2). GFAP, BLBP, and SOX2 (Stage 1) showed the lesser fall (no significant decline) in postnatal expression in comparison to other stem cell marker genes. BLBP showed significant decline only when prenatal expression was compared to adult (Fig. 1, S1 (b.2)). SOX2 showed significant decline only when prenatal expression compared to adolescence and adult age group (Fig. 1, S1 (e.2, 3)). Uniquely, the postnatal sharp downregulation of KI67 and TBR2 continued across advancing age groups, reached near baseline until adolescent age (Fig. 2).

**Figure 2.**
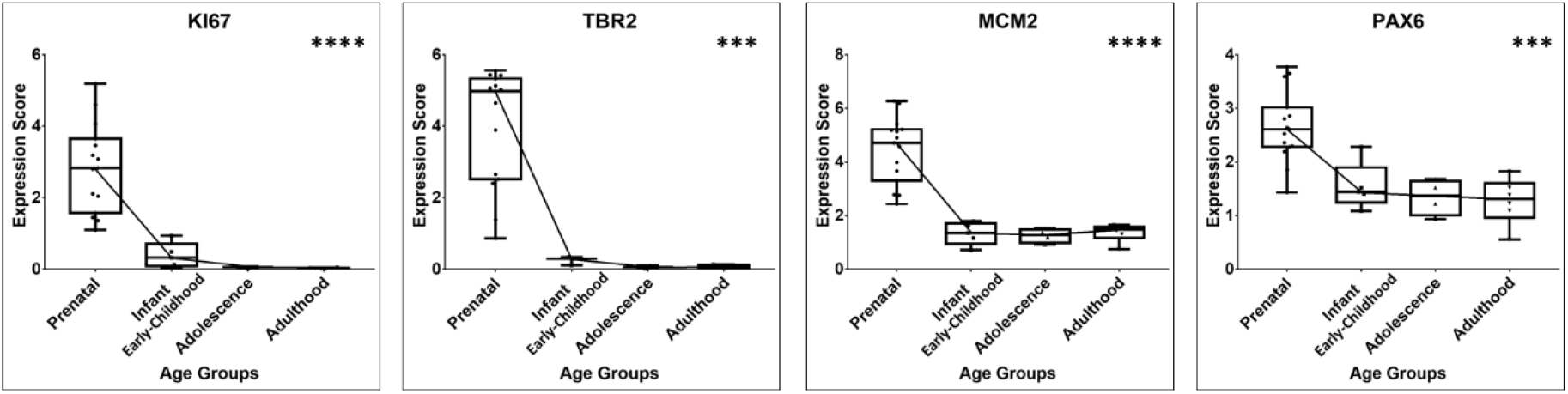
Box plot presentation of Neurogenesis Stage 2 specific gene expression scores (Statistical significance tested using Kruskal Wallis test, p values: *≤0.05, **≤0.01, ***≤0.001, ****≤0.0001).

A post infant/early childhood near maintained expression of immature granule cell markers (NEUROD1, DCX, PSA NCAM)—Stage 3 (Fig. 3), and maintained expression of post mitotic early maturation markers (STMN2, SEMA3C, CALB2, TUBB3) was noted, Stage 4 (Fig. 4). DCX expression showed an increase in level of significance between infant/early childhood to adulthood group (when compared with prenatal values) (Fig. S3 (b.1, 3)).

**Figure 3.**
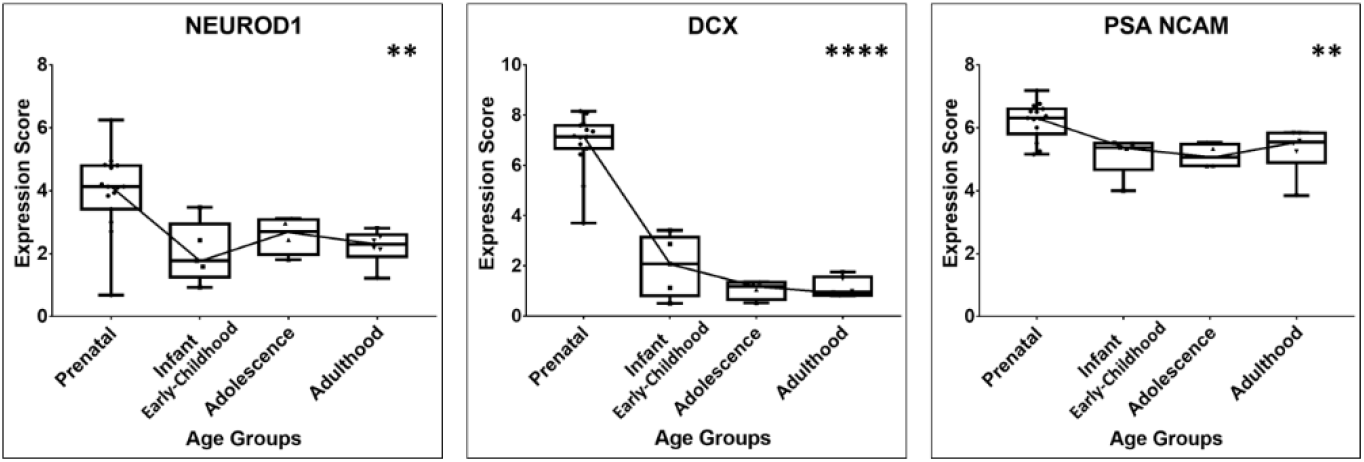
Box plot presentation of Neurogenesis Stage 3 specific gene expression scores (Statistical significance tested using Kruskal Wallis test, p values: *≤0.05, **≤0.01, ***≤0.001, ****≤0.0001.).

**Figure 4.**
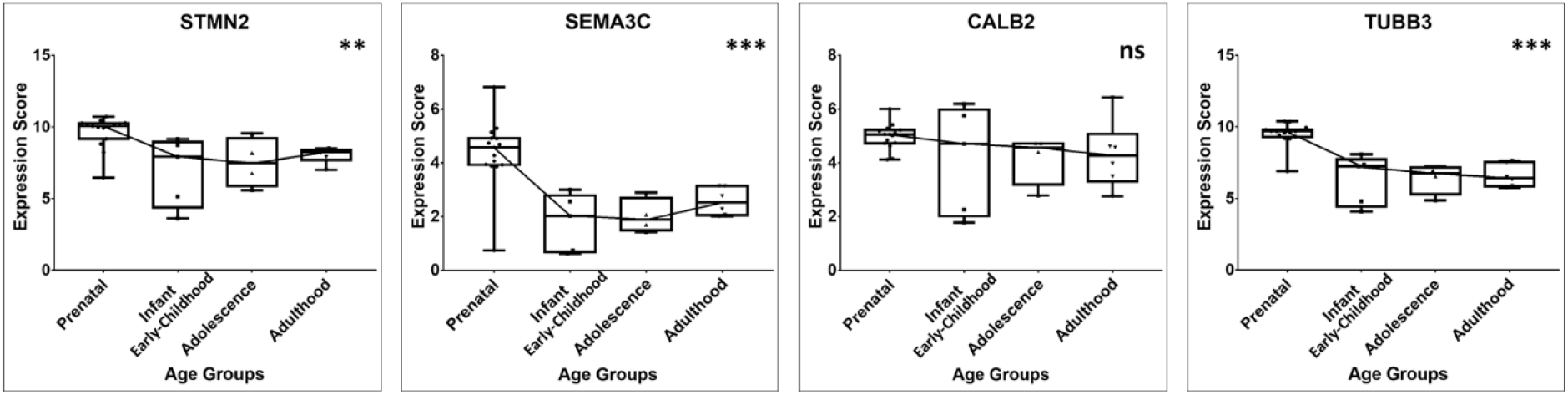
Box plot presentation of Neurogenesis Stage 4 specific gene expression scores (Statistical significance tested using Kruskal Wallis test, p values: *≤0.05, **≤0.01, ***≤0.001, ****≤0.0001, ns=non-significant).

A prenatal to infant/early childhood no difference (though insignificant, the trend of increase for NEUN and CALB1 and in opposite the decreasing trend is observed for MAP2) and thereafter maintained expression of the post mitotic late maturation markers (MAP2, CALB1, NEUN) was noted, Stage 5 (Fig. 5). Uniquely, NEUN showed gain in significance, i.e., increase in its expression is noted in adulthood compared to the prenatal age group. Fig S5 (c.3)).

**Figure 5.**
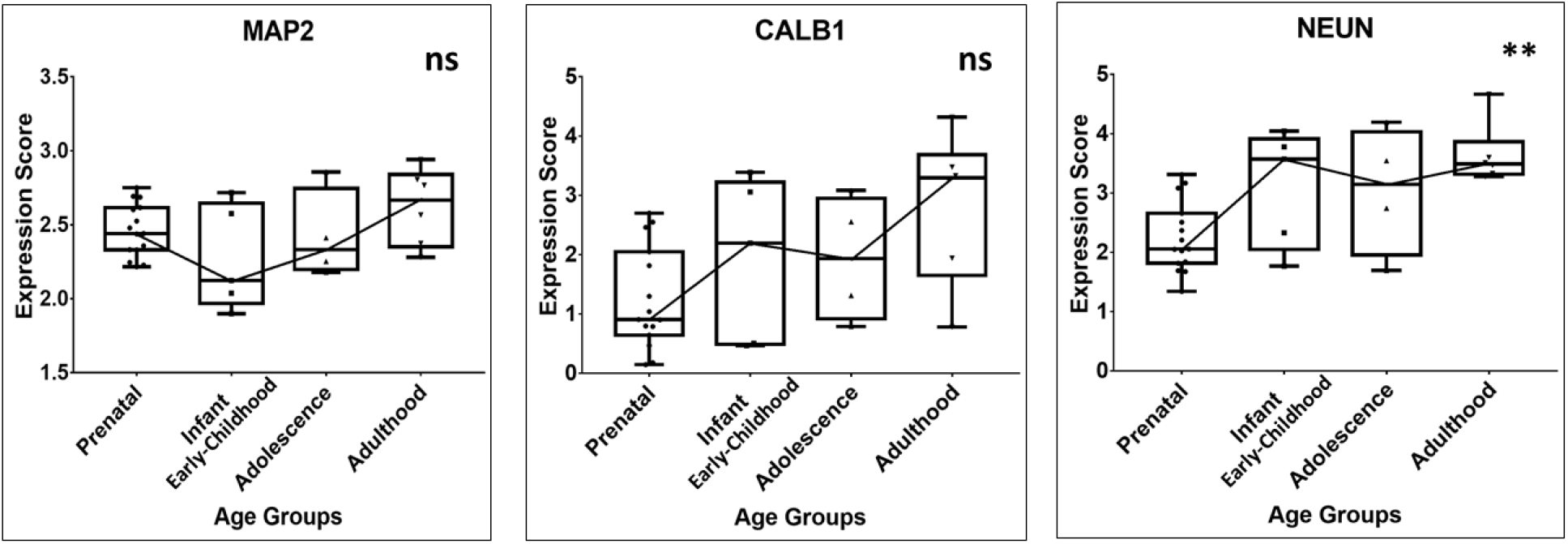
Box plot presentation of Neurogenesis Stage 5 specific gene expression scores (Statistical significance tested using Kruskal Wallis test, p values: *≤0.05, **≤0.01, ***≤0.001, ****≤0.0001, ns=non-significant).

Amongst stage 1-5, no markers showed any significant change in expression among postnatal age groups except, SOX4 (Stage 1) which showed significant decline between infant/early childhood and adult age (Fig. S1 (f.5)). There existed no significant differences in the expression across age groups (all MWU comparisons) for GFAP (Stage 1), PSA-NCAM (Stage 3) CALB2 (Stage 4), and CALB1, MAP2 (Stage 5).

### Gliogenesis markers

OLIG 2 and VIMENTIN showed significant decline in their gene expression from prenatal to postnatal transition, S100beta (S100B) showed significant increase (~3 folds) during this transition (Fig. 6, S6, Table S1&2). All the gliogenesis markers showed no significant differences in expression between the postnatal age groups, though trend of curve declined from infant/early childhood to adult with increase in level of significance (when compared to prenatal values) (Fig. S6).

**Figure 6.**
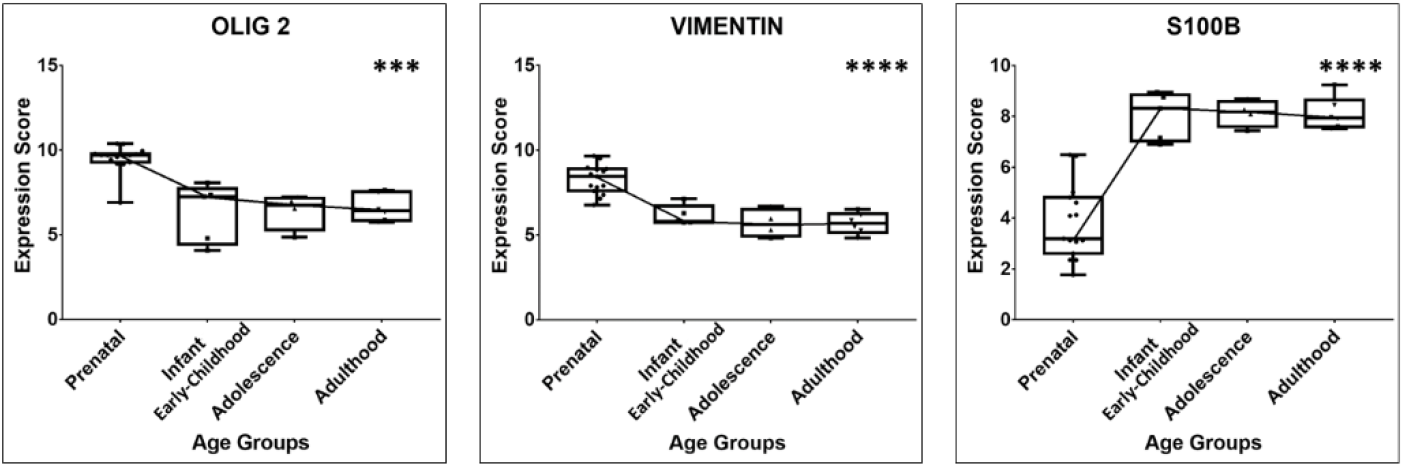
Box plot presentation of Gliogenesis gene expression scores (Statistical significance tested using Kruskal Wallis test, p values: *≤0.05, **≤0.01, ***≤0.001, ****≤0.0001).

### Apoptotic markers

BAX and TP53 showed significant decline in expression between prenatal to infant/early childhood (Fig. 7&S7). For both the markers declining trend continued across age groups with no statistically significant differences for comparisons between close groups (Fig. 7&S7).

**Figure 7.**
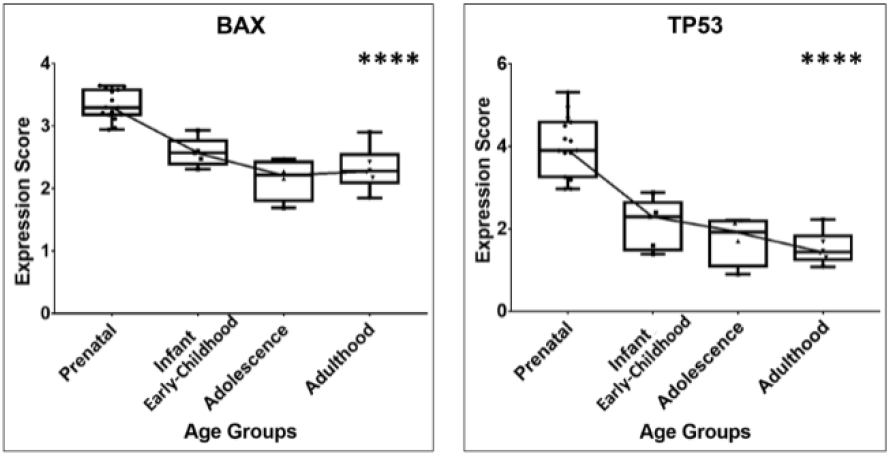
Box plot presentation of Apoptotic gene expression scores (Statistical significance tested using Kruskal Wallis test, p values: *≤0.05, **≤0.01, ***≤0.001, ****≤0.0001).

## Discussion

Study of the hippocampal developmental transcriptome gives a glimpse of the events happening in the neurogenic niche. We studied hippocampal developmental transcriptome in chronological detail and looked for the stage specific expression of neurogenesis signature, gliogenesis, and apoptotic markers. This led to imperative observations which may pave way for future studies to resolve the intense controversy surrounding AHN.

Our results (Table S1&2, Fig. 1-7, S1-7) showed that there are contrasting differences in the expressions of the stage specific hippocampal neurogenesis markers when we compare the developmentally advancing age groups. Plausibly, in case of no neurogenesis, all neurogenesis signature markers, irrespective to the stages, should have shown significant downfall in expression with the advancement of the age. The analyses of expression patterns of the studied marker genes indicate that though stem cells and progenitor cell forms (Stage 1&2) may be decreasing (though negligibly between two close age groups), the total number of immature granule cells (Stage 3) and post mitotic forms as early mature and mature granule cells (Stage 4&5)—is perhaps maintained from prenatal/childhood to adult stages of the human life cycle. Similar conclusions were derived from the study by Boldrini et al., in age range 14-79 year, using immunohistological methods, providing evidence that new neurons keep forming in the neurogenic niche of the adult human hippocampus also.

A postnatal sharp fall in the expression of most of the stem cell markers (NESTIN, SOX1, and SOX4) (Fig. 1), and thereafter decreasing trend is indicative of a considerable postnatal downfall in production of NSCs. A prenatal-to-childhood sharp decrease in the NSCs has been a common observation in the human developmental studies (3–6). Boldrini et al., noted a decrease of NSCs with aging in age group 14-79 year. Our data showed a trend that, a post childhood decline is very slow, and near maintained between adolescent and adult age groups. GFAP, BLBP and SOX 2 showed a relatively lesser downregulation (only childhood to adolescence: SOX2, and only adolescence to adult: BLBP showed significant decline, no significant difference from prenatal to adult is noted for GFAP) in comparison to the other stem cell markers, this could be explained by the fact that other than NSCs, they also express in glial lineage cells in adult brain.

A post childhood near maintained expression of immature granule cell markers (DCX, PSA NCAM, NEUROD1 didn’t show a significant difference between age groups)—Stage 3 (Fig. 3), and maintained expression of the post mitotic early maturation markers (SEMA3C, STMN2, TUBB3, CALB2: Calretinin is a calcium binding molecule, which is expressed up to 2-3 weeks in newly born neurons)—Stage 4 (Fig. 4), hint for the continued formation of neuronal lineage cells and their further differentiation into immature and early mature neurons.

The expression patterns (post childhood maintained or increased insignificantly between two terminal age groups) of the post mitotic marker genes (Stage 5) (Fig. 5) NEUN, and calbindin (CALB1)—another calcium marker which follows the transient expression of calretinin (CALB2) (after 2-3 weeks in young neurons), and MAP2, further add up to the evidence for the continued maturation of newborn neurons taking place in adult hippocampus.

Though, a postnatal steeply decreasing trend of the transiently amplifying progenitor cell marker genes (Stage 2, Fig. 2), especially KI67, TBR2 indicates that, constrained by the sharp decline and progressively low rate of mitosis post infant/early childhood age, only a minimal formation of new cells (in comparison to prenatal values) at any point of time would be possible until adult age. Sorrells et al found that, at 22 gestational weeks, KI67^+^ cells were less than 100/mm^2^ which decreased sharply during the first year of life, and there were rare instances of KI67^+^ cells in the DG of a 35-year-old individual. In same study, Sorrells et al, performed a comparative developmental transcriptomic analysis (between human and macaque monkey) of KI67 gene, and found a similar pattern, as we show in our study (3). Boldrini et al found KI67^+^ cells (without distinction of non-neuronal or endothelial cells) were in the order of 10^4^/ DG region (anterior, mid, and posterior) and unchanged between 14 and 79 years of age (6). Tobin et al, reported NESTIN^+^ KI67^+^ (Stage 2) cells 1,437.8 ± 274.2 for entire DG in the individuals between 79 and 99 years (8).

In this study, we are not able to provide a clear estimate of the new cells count which can be formed in adult age, but in view of the steep decline of certain Stage 2 markers—KI67 (which denotes dividing cells), and TBR2, we assume that, it has to be negligible in comparison to the prenatal values. Boldrini et al, 2018, and Moreno-Jiménez et al, 2019 showed number of newly formed neurons as many as 3 thousands/DG region, and 30 (± 2.5) thousands/mm^3^ of DG respectively (6,7). Tobin et al found 127,342 ± 28,864 DCX^+^ cells (neuroblasts and immature neurons) for entire DG (8). Though, data from all three studies are not exactly comparable due to difference in the target age group, they certainly present strong evidence for AHN. The variation in the number of newborn cells may also have been contributed by differences in PMI, fixation time, and tissue processing for each study. Unfortunately, all three of the studies didn’t involve prenatal age participants, so it can’t be made out from their studies, in what fraction, neurogenesis was reduced in postnatal age groups (when compared to the prenatal values). Sorrels et al, who studied both prenatal and adult age group showed 1600 ± 800/mm^2^ of DG in prental brain, with steeply decresing trend in postnatal age groups, almost nil in adult age (3). Though, the tissue fixation and processing methods Sorrels et al used, might have caused low show of neuronal lineage cells across all ages in their study, as was also suggested by Moreno-Jiménez et al, 2019, who used improvised immunohistological methods (7). A study in prenatal-to-adult age group using tightly controlled conditions and state-of-the-art tissue processing methods, as has been suggested by Moreno-Jiménez et al (7), may bring some more clarity on age group specific formation of new neurons.

Uniquely, a continued postnatal decrease in apoptotic markers (BAX, and TP53) with insignificant differences between advancing age groups, gives clear indications that there is no increased cell death among the advancing age groups hence total number of the mature neurons may be maintained with aging as was claimed by Boldrini et al (for 14-79 year age group) (6). Surprisingly, apoptotic markers were scarcely examined in any histological study in this regard. A maintained total number of hippocampal neurons further hints for a minimal but continued neurogenesis which would be just sufficient to replenish the average loss of neurons with aging. A minimal but continued neurogenesis in adult human hippocampus is plausible in view of the available studies which support the presence of stem cell pool in the neurogenic niche which can be engaged in the maintenance of total number of neurons (6,22).

In contrast to the neuron lineage cell markers, expression patterns of the gliogenesis markers—VIMENTIN (a pan astrocytic marker), S100B (a post mitotic astrocytic marker), and OLIG 2 (an oligodenderocyte lineage marker), suggest that a significant part of the new born cells in the neurogenic niche in adult human hippocampus may be glial cells.

Interpretations of our data advise against the conclusion made by Dennis et al, Sorrels et al, and Cipriani et al (3–5), that there is absolute post childhood seize in formation of new neurons, though we support the notion that a significant proportion of the new formed cells in the neurogenic niche may be the glial cells. We conclude that an active but minimal hippocampal neurogenesis is possibly continued in adult human.

Limitations of this study are: (i) neural tissue used for transcriptomic analysis in original data source were not exclusively taken from neurogenic niche in DG, homogenate from whole hippocampus is used, (ii) an uneven and relatively smaller sample size for some age groups, (iii) additionally, many of the neurogenesis markers we studied are known to express in more than one consecutive developmental stages, so their exact demarcation respective to the stages was not feasible.

Taking a homogenate from whole hippocampus might have some impact on the absolute expression value of neurogenesis markers. Though, that couldn’t have affected relative expression of studied markers across age group hence would have made little impact on our analysis. We believe that influences of uneven and relatively smaller sample size have been largely cancelled by selecting suitable non parametric tests to compute differential expression of the genes. Further integrative studies design across age group combining the state of art immunohistological and transcriptomic methods may provide a clearer view on the continuity of AHN.

## Supporting information

Significance statement

## Acknowledgments

Data for this study was collected from the BrainSpan Atlas of the Developing Human Brain. The findings of this study were first presented at the Annual Meeting of Society for Neuroscience (SFN), 3^rd^-7^th^ November, 2018, San Diego, USA.

## Author (s) contributions

AK conceived, and AK and VP designed the study. AK, VP analysed the data and wrote the first draft. AK, VP, MF, PK, CK, HS, SG revised the first draft. All authors have consented for the submission of the final draft.

## Supplementary Files

### Results

**Figure S1.**
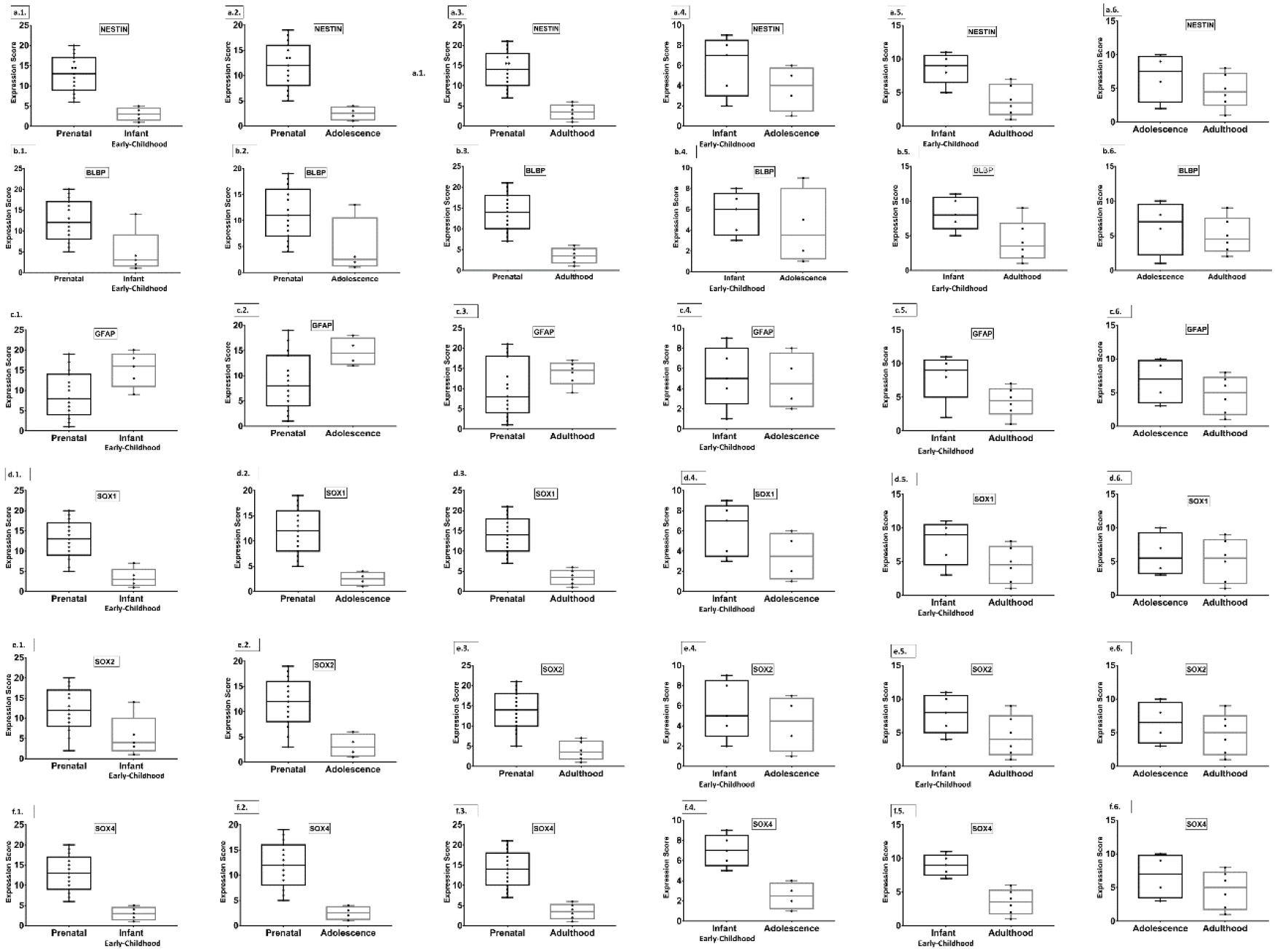
Box plot presentation of Neurogenesis Stage 1 specific gene expression scores (Statistical significance tested using Mann Whitney U test).

**Figure S2.**
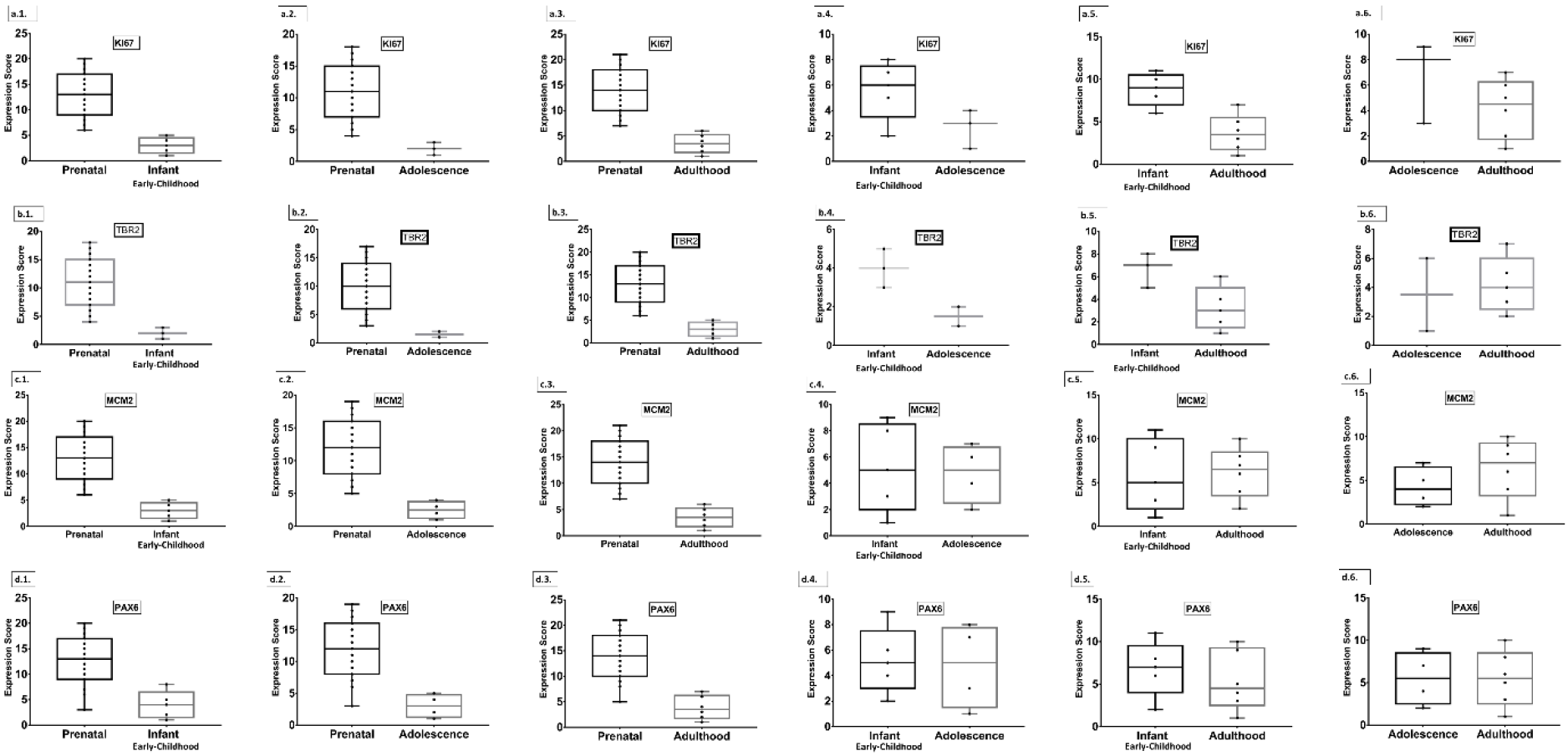
Box plot presentation of Neurogenesis Stage 2 specific gene expression scores (Statistical significance tested using Mann Whitney U test).

**Figure S3.**
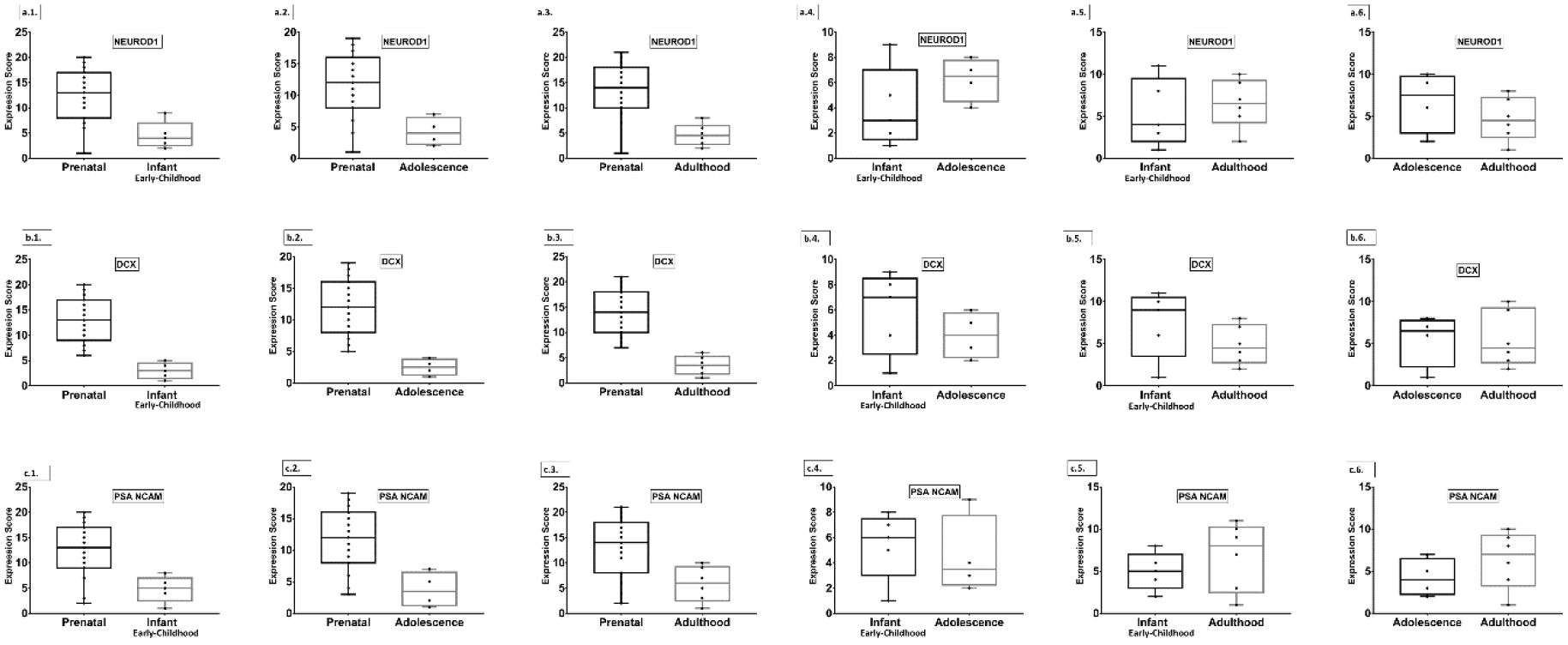
Box plot presentation of Neurogenesis Stage 3 specific gene expression scores (Statistical significance tested using Mann Whitney U test).

**Figure S4.**
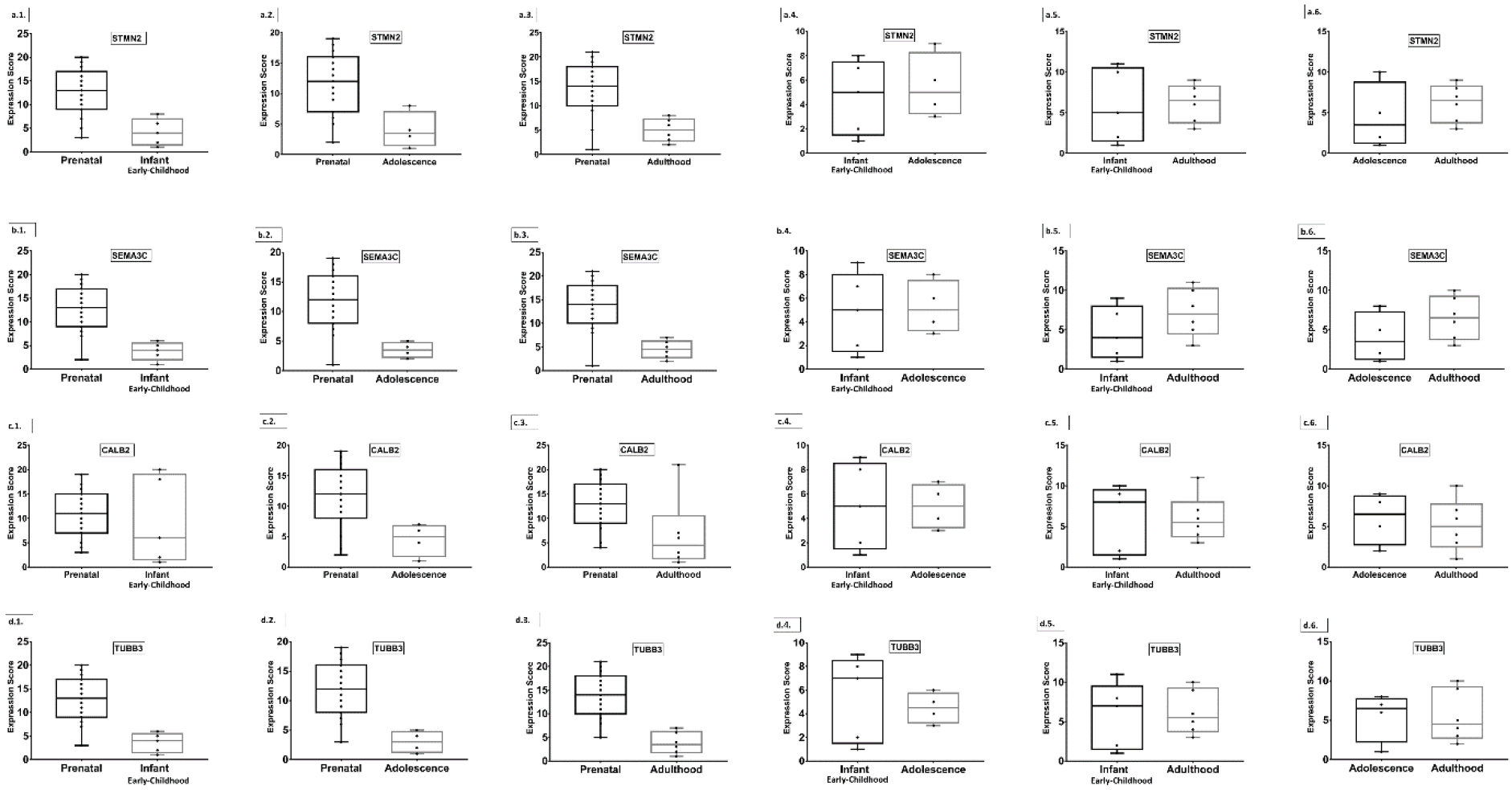
Box plot presentation of Neurogenesis Stage 4 specific gene expression scores (Statistical significance tested using Mann Whitney U test).

**Figure S5.**
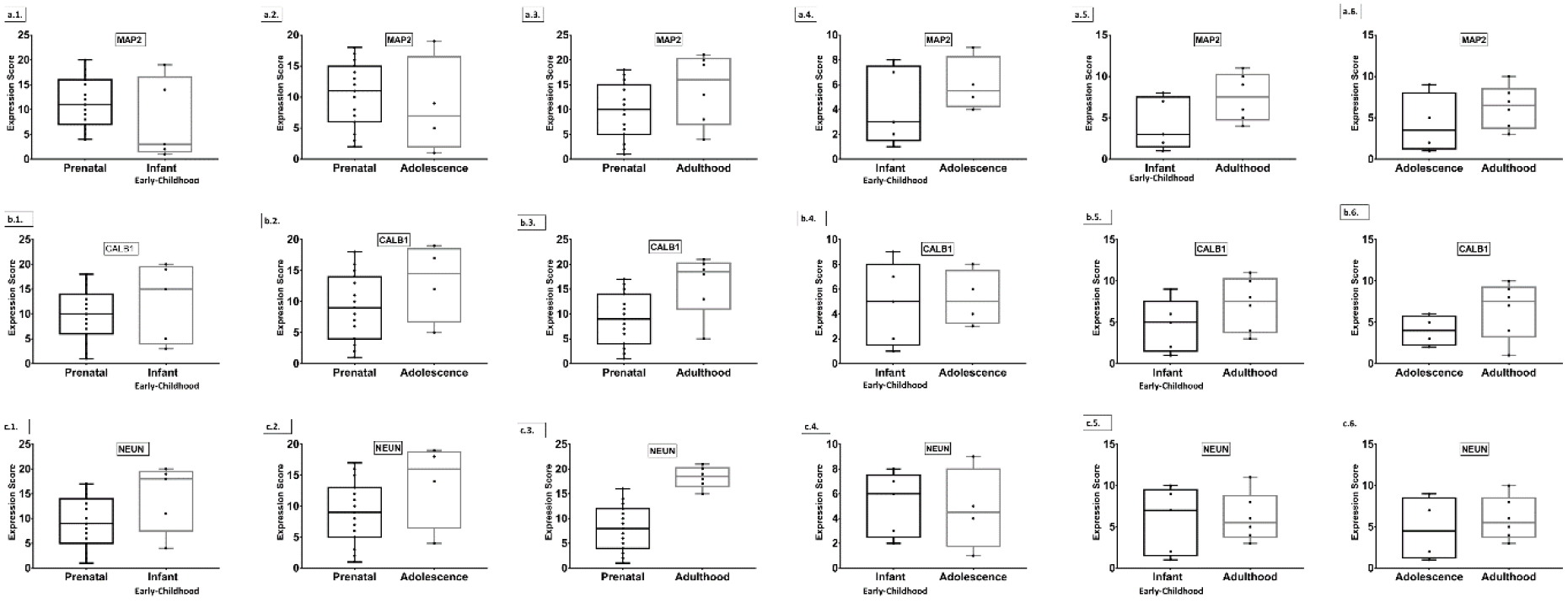
Box plot presentation of Neurogenesis Stage 5 specific gene expression scores (Statistical significance tested using Mann Whitney U test).

**Figure S6.**
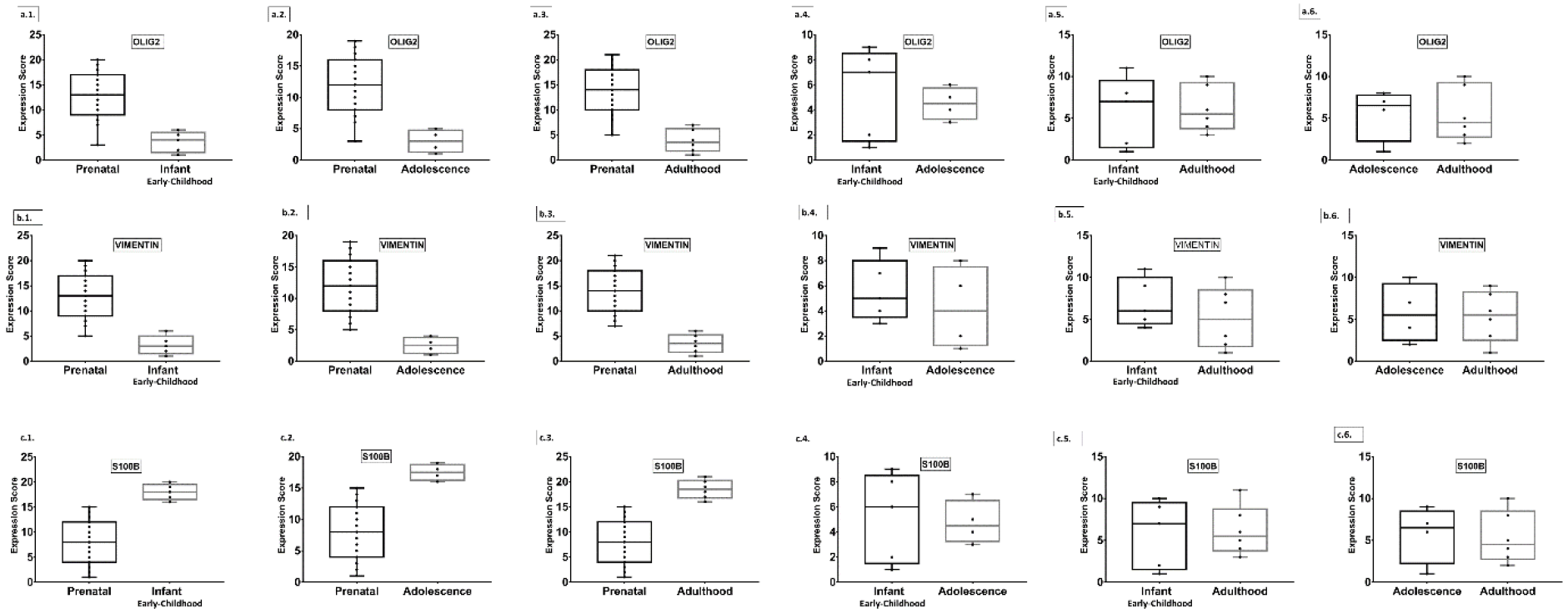
Box plot presentation of Gliogenic gene expression scores (Statistical significance tested using Mann Whitney U test).

**Figure S7.**
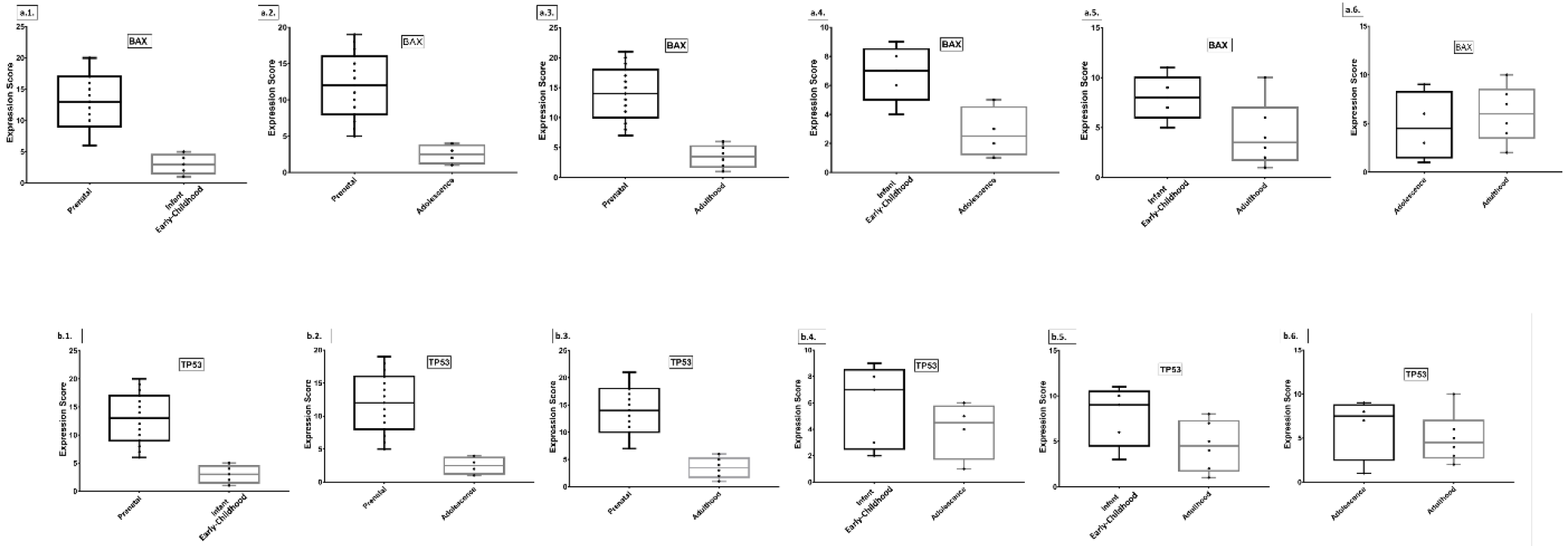
Box plot presentation of Apoptotic gene expression scores (Statistical significance tested using Mann Whitney U test).

**Table S1:** Differential expression of Neurogenesis, gliogenesis, apoptotic markers across developing age groups. (Statistical significance tested using Kruskal Wallis test (KW)). *[Median_P(Prenatal), Median_I(Infant/Early-Childhood), Median_Ae(Adolescence), Median_Ad(Adulthood)].*

| Category | Genes | P value | KW | n | *Median_P* | *Median_I* | *Median_Ae* | *Median_Ad* |
| --- | --- | --- | --- | --- | --- | --- | --- | --- |
| **Neurogenesis Markers** |  |  |  |  |  |  |  |  |
| **Stage 1:** | NESTIN | <0.0001 | 23.31 | 30 | 6.17 | 4.418 | 3.38775 | 2.921 |
|  | BLBP | 0.0007 | 17.13 | 30 | 6.4209 | 5.4084 | 4.75115 | 4.47175 |
|  | GFAP | 0.0496 | 7.832 | 30 | 0.6461 | 1.86 | 1.7084 | 1.5429 |
|  | SOX1 | <0.0001 | 22.06 | 30 | 4.0465 | 1.6384 | 1.1075 | 1.01615 |
|  | SOX2 | 0.0007 | 17.01 | 30 | 6.664 | 5.0575 | 4.8076 | 4.1527 |
|  | SOX4 | <0.0001 | 24.33 | 30 | 7.452 | 2.9735 | 1.9225 | 1.5461 |
| **Stage 2:** | KI67 | <0.0001 | 22.88 | 29 | 2.8357 | 0.3222 | 0.0575 | 0.0295 |
|  | TBR2 | 0.0004 | 18.12 | 25 | 4.9813 | 0.2954 | 0.0603 | 0.0751 |
|  | MCM2 | <0.0001 | 21.97 | 30 | 4.7084 | 1.3511 | 1.2746 | 1.4784 |
|  | PAX6 | 0.0003 | 18.63 | 30 | 2.6077 | 1.4414 | 1.37035 | 1.31075 |
| **Stage 3:** | NEUROD1 | 0.0021 | 14.66 | 30 | 4.1275 | 1.777 | 2.6986 | 2.29665 |
|  | DCX | <0.0001 | 22.24 | 30 | 7.1393 | 2.0775 | 1.18225 | 0.96115 |
|  | PSA NCAM | 0.0047 | 12.95 | 30 | 6.3157 | 5.3719 | 5.06755 | 5.54735 |
| **Stage 4:** | STMN2 | 0.0023 | 14.52 | 30 | 10.066 | 7.9271 | 7.4753 | 8.2299 |
|  | SEMA3C | 0.0006 | 17.34 | 30 | 4.5705 | 2.0226 | 1.88495 | 2.52845 |
|  | CALB2 | 0.1377 | 5.517 | 30 | 5.0564 | 4.7054 | 4.564 | 4.2732 |
|  | TUBB3 | 0.0003 | 19.17 | 30 | 9.6975 | 7.2513 | 6.74995 | 6.4186 |
| **Stage 5:** | MAP2 | 0.2296 | 4.312 | 30 | 2.4405 | 2.1235 | 2.33185 | 2.66665 |
|  | CALB1 | 0.0873 | 6.56 | 30 | 0.9087 | 2.1942 | 1.9338 | 3.2945 |
|  | NEUN | 0.0088 | 11.63 | 30 | 2.0574 | 3.573 | 3.144 | 3.48925 |
| **Gliogenesis Markers** | OLIG 2 | 0.0003 | 19.17 | 30 | 9.6975 | 7.2513 | 6.74995 | 6.4186 |
|  | VIMENTIN | <0.0001 | 21.69 | 30 | 8.4493 | 5.8152 | 5.6258 | 5.68415 |
|  | S100B | <0.0001 | 21.77 | 30 | 3.1896 | 8.3201 | 8.17725 | 7.94155 |
| **Apoptotic Markers** | BAX | <0.0001 | 23.28 | 30 | 3.2914 | 2.5705 | 2.2137 | 2.2757 |
|  | TP53 | <0.0001 | 22.52 | 30 | 3.9032 | 2.2932 | 1.92155 | 1.43985 |

**Table S2:** Differential expression of Neurogenesis, gliogenesis, apoptotic markers between close developing age groups. (Statistical significance tested using Mann Whitney U test, Median expressions). *[P=Prenatal, I=Infant/Early-Childhood, Ae=Adolescence, Ad=Adulthood].* *(The Medians for age specific expressions of all genes compared in Mann Whitney U test are same as in **Table S1** (refer that). Level of significance set at p≤0.05.)*

| Category | Genes | *U stats and Level of significance scores in successive orders (U value, p-value)* |  |  |  |  |  |
| --- | --- | --- | --- | --- | --- | --- | --- |
| Neurogenesis Markers |  | *P vs I* | *P vs Ae* | *P vs Ad* | *I vs Ae* | *I vs Ad* | *Ae vs Ad* |
| Stage 1: | NESTIN | 0, 0.0001 | 0, 0.0005 | 0, 0.00001 | 5, 0.28 | 2, 0.017 | 7, 0.35 |
|  | BLBP | 9, 0.010 | 9, 0.036 | 0, 0.000036 | 7, 0.55 | 4, 0.05 | 9, 0.60 |
|  | GFAP | 14, 0.041 | 11, 0.062 | 28, 0.205 | 9, 0.90 | 5, 0.08 | 7, 0.35 |
|  | SOX1 | 2, 0.00051 | 0, 0.00051 | 0, 0.000036 | 4, 0.19 | 6, 0.12 | 10, 0.76 |
|  | SOX2 | 13, 0.032 | 3, 0.0036 | 2, 0.00014 | 7, 0.55 | 6, 0.125 | 8, 0.47 |
|  | SOX4 | 0, 0.00012 | 0, 0.00051 | 0, 0.000036 | 0, 0.015 | 0, 0.004 | 7, 0.35 |
| Stage 2: | KI67 | 0, 0.0001 | 0, 0.002 | 0, 0.00003 | 2, 0.14 | 1, 0.008 | 4, 0.26 |
|  | TBR2 | 0, 0.0024 | 0, 0.014 | 0, 0.00012 | 0, 0.2 | 1, 0.071 | 4, 0.85 |
|  | MCM2 | 0, 0.0001 | 0, 0.000515 | 0, 0.000036 | 9, 0.90 | 14, 0.93 | 7, 0.35 |
|  | PAX6 | 5, 0.0024 | 2, 0.0020 | 2, 0.00014 | 9, 0.90 | 11, 0.53 | 12, >0.99 |
| Stage 3: | NEUROD1 | 8, 0.0077 | 7, 0.019 | 7, 0.0016 | 5, 0.28 | 12, 0.66 | 7, 0.35 |
|  | DCX | 0, 0.00012 | 0, 0.00051 | 0, 0.00003 | 6, 0.41 | 8, 0.24 | 12, >0.99 |
|  | PSA NCAM | 9, 0.010 | 5, 0.0092 | 14, 0.014 | 8, 0.73 | 10, 0.42 | 7, 0.35 |
| Stage 4: | STMN2 | 0, 0.0037 | 6, 0.013 | 9, 0.0033 | 8, 0.73 | 14, 0.93 | 8, 0.47 |
|  | SEMA3C | 4, 0.0015 | 4, 0.0061 | 6, 0.001 | 9, 0.90476 | 8, 0.24 | 6, 0.25 |
|  | CALB2 | 32, 0.67 | 8, 0.027 | 19, 0.04 | 10, >0.99 | 15, >0.99 | 10, 0.76 |
|  | TUBB3 | 3, 0.0009 | 2, 0.0020 | 2, 0.00014 | 8, 0.73 | 14, 0.93 | 12, >0.99 |
| Stage 5: | MAP2 | 24, 0.26 | 24, 0.59 | 26, 0.15 | 6, 0.41 | 6, 0.12 | 7, 0.35 |
|  | CALB1 | 28, 0.44 | 17, 0.22 | 15, 0.018 | 9, 0.90 | 8, 0.24 | 6, 0.25 |
|  | NEUN | 18, 0.09 | 15, 0.15 | 1, 0.000073 | 9, 0.90 | 14, 0.93 | 9, 0.60 |
| Gliogenesis Markers | OLIG 2 | 3, 0.0009 | 2, 0.002 | 2, 0.0001 | 8, 0.73 | 14, 0.93 | 12, >0.9999 |
|  | VIMENTIN | 1, 0.000257998 | 0, 0.0005 | 0, 0.00003 | 7, 0.55 | 10, 0.42 | 11, 0.91 |
|  | S100B | 0, 0.00012 | 0, 0.00051 | 0, 0.000036 | 9, 0.90 | 14, 0.93 | 11, 0.91 |
| Apoptotic Markers | BAX | 0, 0.0001 | 0, 0.0005 | 0, 0.00003 | 1, 0.031 | 5, 0.082 | 9, 0.60 |
|  | TP53 | 0, 0.0001 | 0, 0.0005 | 0, 0.00003 | 6, 0.41 | 6, 0.12 | 9, 0.60 |

