## Supplementary material for "Transcriptomic Analysis of the Neurogenesis Signature suggests Continued but Minimal Neurogenesis in the Adult Human Hippocampus": Significance statement

Immunohistological investigations have given extremely contradictory results about continued hippocampal neurogenesis in adult humans. Studies are scarce which explored hippocampal neurogenesis across advancing age groups; perhaps, it has been due to difficulty in the procurement of the postmortem brain tissue in optimized laboratory conditions. Developmental human brain transcriptome, which is freely available on the Allen Brain Atlas website provides an excellent opportunity to look for the expression status of the neurogenesis markers in the hippocampus across the advancing age groups. Also, until now, not much stress has been given to check expressions of the gliogenesis and apoptotic markers in this paradigm, which could provide essential inferences. We performed a transcriptomic analysis of the neurogenesis signature supplemented with the gliogenesis and apoptotic markers in the hippocampal data in context to the developmental stages across age groups. Results of this study have led to a conclusion which stands between the two contrasting views from the immunohistological studies—which has come either in absolute denial or significant approval for the hippocampal neurogenesis in the adult humans. We propose that a continued but minimal neurogenesis may be the original feature in the adult human hippocampus.
